# Tracking the Morphological Evolution of Neuronal Dendrites by First-Passage Analysis

**DOI:** 10.1101/2024.10.13.618072

**Authors:** Fabian H. Kreten, Barbara A. Niemeyer, Ludger Santen, Reza Shaebani

## Abstract

A high degree of structural complexity arises in dynamic neuronal dendrites due to extensive branching patterns and diverse spine morphologies, which enable the nervous system to adjust function, construct complex input pathways and thereby enhance the computational power of the system. Owing to the determinant role of dendrite morphology in the functionality of the nervous system, recognition of pathological changes due to neurodegenerative disorders is of crucial importance. We show that the statistical analysis of a temporary signal generated by cargos that have diffusively passed through the complex dendritic structure yields vital information about dendrite morphology. As a feasible scenario, we propose engineering mRNA-carrying multilamellar liposomes to diffusively reach the soma and release mRNAs, which are translated into a specific protein upon encountering ribosomes. The concentration of this protein over a large population of neurons can be externally measured, as a detectable temporary signal. Using a stochastic coarse-grained approach for first-passage through dendrites, we connect the key morphological properties affected by neurodegenerative diseases—including the density and size of spines, the extent of the tree, and the segmental increase of dendrite diameter towards soma—to the characteristics of the evolving signal. Thus, we establish a direct link between the dendrite morphology and the statistical characteristics of the detectable signal. Our approach provides a fast noninvasive measurement technique to indirectly extract vital information about the morphological evolution of dendrites in the course of neurodegenerative disease progression.

## INTRODUCTION

The elaborate branching morphology of neuronal dendrites in advanced nervous systems allows the neuron to interact simultaneously with several neighbors through their axon terminals, leading to a complex network of signaling pathways [1, 2]. The diverse functions of dendritic trees are reflected in the broad variation of their architecture in different neuronal types and regions. The complex behavior of higher animals has also been attributed to the presence of small membranous protrusions called dendritic spines [3–6]. Functional synapses, as the basic computational units of signal transmission, consist of the presynaptic release site and dendritic protrusions, harbouring the signal recognition and transmission units. Spines play a vital role in neural functions such as cognition, memory, and learning [7–11] and often serve as the recipients of excitatory and inhibitory inputs in the mammalian brain and undergo dynamic structural changes regulated by neuronal activity [12, 13]. The morphology of spines plays a crucial role as, for example, the shape and size of spine head and neck determine the number of postsynaptic receptors and the generated synaptic current [8] and control the electrical and biochemical isolation of the spine from the dendrite shaft [14, 15].

Aging and several neurodegenerative diseases—e.g. fragile X and Down’s syndromes, Alzheimer’s disease, schizophrenia, and autism spectrum disorders—significantly influence the function of the nervous system by altering the morphology of dendrites [16–20]. These alterations occur in the overall extent of dendritic trees, the population of branches, the thickness and curvature of dendrite shafts, and the density, morphology, and spatial distribution of spines [21–34]. On the other hand, reversal of morphological changes upon treatment has also been reported [35, 36]. Despite the crucial importance of monitoring the structural evolution of dendritic trees and spines to diagnose and predict neurodegenerative diseases and to monitor success of potential treatments, noninvasive imaging of neuronal dendrites is currently infeasible. It is even highly challenging to collect statistically adequate structural information from direct invasive imaging due to technical limitations: Although image analysis techniques for 3D reconstruction of dendrites have been improved in recent years [37, 38], a high resolution image can be achieved by electron microscopy which is a very laborious technique and practically inappropriate for spatially large-scale investigations. Nevertheless, there exist powerful noninvasive techniques which allow real-time tracking of brain activities, ranging from electro- and magneto-encephalography for electric and magnetic field detection [39, 40] to nuclear magnetic resonance spectroscopy, positron emission tomography, and magnetic resonance imaging (MRI) for measuring the concentration of (neuro-)chemicals [41–44].

In the absence of a direct efficient method for unraveling the microscopic morphology of spines and dendrites, we alternatively develop a fast noninvasive technique—based on processing an externally detectable signal generated by a large population of neurons—to indirectly obtain essential structural information. A possible realization of such a signal can be the concentration of an expressed protein by many neurons in the brain region of interest, monitored e.g. by the MRI technique. We discuss how mRNA-carrying multilamellar liposomes can reach a desired region of brain tissue using brain-targeted drug delivery techniques, pass through the complex dendrite structures to reach the soma, and release their mRNAs which encodes a specific protein upon encountering ribosomes (alternatively, liposomes could carry siRNA molecules to silence RNA and decrease a specific protein concentration). We develop a framework that bridges the diverse morphological characteristics of dendrites and the statistical properties of the detectable evolving signal (linked to the first-passage times of passing through the dendrite structure). Our technique enables systematic monitoring of neurodegenerative disease or treatment progression for individual patients.

### COARSE-GRAINED DENDRITE MODEL

To model the structure of dendrites, we adopt a mesoscopic perspective and consider a regularly branched tree with *n* generations of junctions. Denoting the average distance between adjacent nodes by *L*, the tree has a linear extent *nL*. Importantly, as we will discuss, our results remain valid under realistic levels of global variation in structural parameters or local irregularities across the tree architecture [45, 46]. By coarse-graining the stochastic transport within dendritic shafts and spines, we study the dynamics of noninteracting particles hopping between the nodes of the tree (see [45] and Fig. 1). Each particle, drawn from an initial reservoir containing *N* particles, enters the tree at a random node indexed by its depth *i* ∈ [0, *n*], where *i* = 0 corresponds to the soma and *i* = *n* to the dead ends. The time of entry is randomly sampled from a geometric distribution with mean *t*_*e*_, consistent with the goal of monitoring transport behavior over a brain region rather than focusing on individual particles. Assuming a uniform probability of arrival across the tree, the entry probability increases exponentially with node depth in a binary tree, yielding an entry-level distribution *Ei* = 2^*i*−1^ */*(2^*n*^−1) for *i*≥1.

**FIG. 1.**
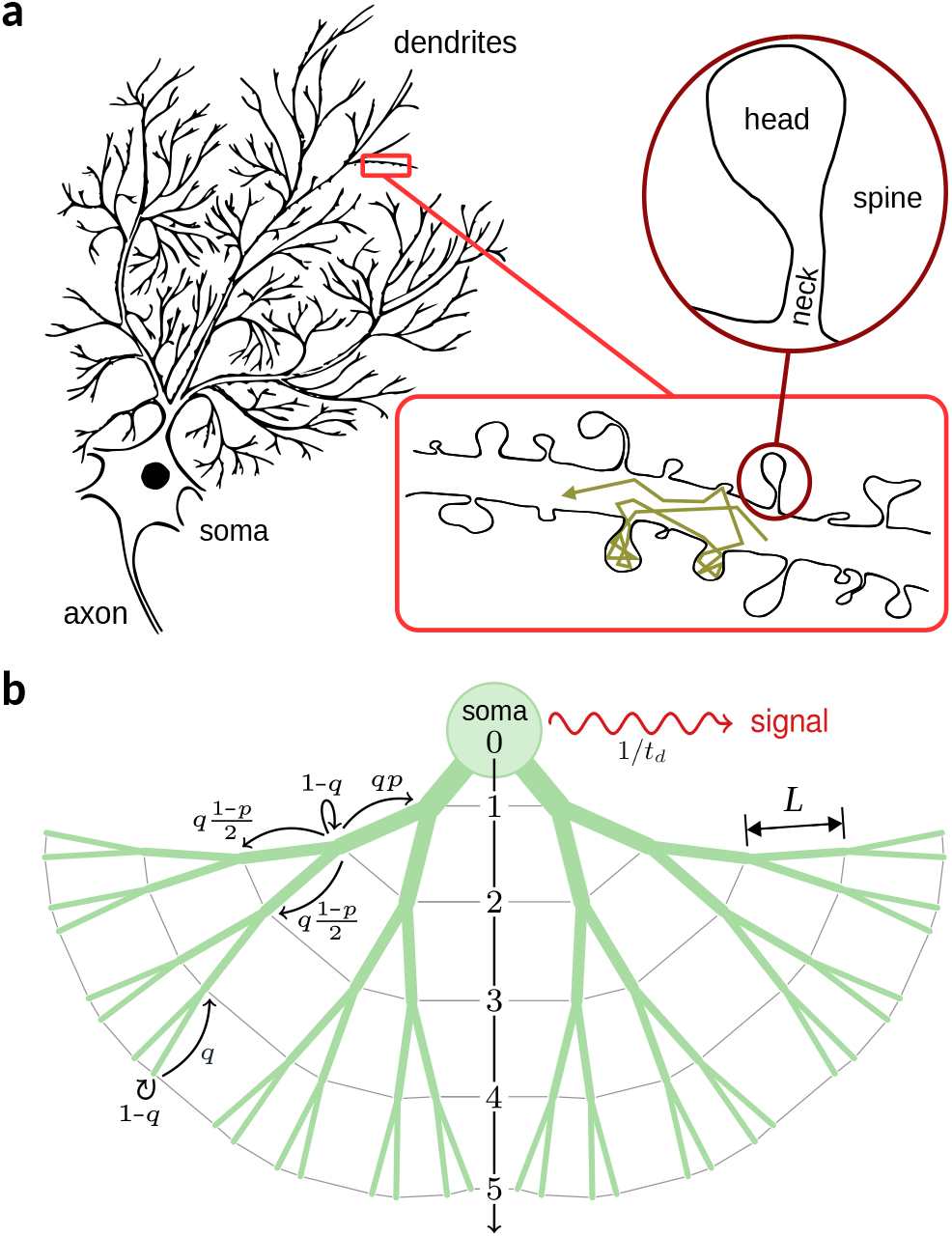
Morphology of neuronal dendrites. (a) Schematic drawing of a neuron. Insets: (lower) A section of a typical dendritic channel. A sample path of a particle is shown; (upper) Structure of a mushroom-like spine. (b) Sketch of our binary tree model. A tree structure with depth *n* = 5 is depicted as an example. The possible choices at junctions or dead-ends are shown with arrows. The coarse-grained probability to reach a neighboring intersect is denoted with *q*, the topological bias *p* represents the segmental increase of dendrite diameter towards the soma, and 1*/t*_*d*_ denotes the pulse emission rate.

At each time step, a particle either hops to a neighboring node with probability *q*, or remains at its current position with probability 1 −*q*. The waiting probability accounts for both stochastic trapping inside dendritic spines and the diffusive delay within the dendrite channel. We assume that the residence probability inside these biochemical cages is depth-independent since the spine number density along the dendrite is known to rapidly saturate after a short distance from the soma [37, 47]. To model the directional preference in tracer particle motion toward the soma or dead ends, we introduce a topological bias parameter *p*, which governs the directional preference of particle movement: transitions toward the soma and toward the dead ends occur with probabilities *p* and 1 −*p*, respectively. Each particle is eventually absorbed at the soma. Upon arrival, it emits a transient pulse after a randomly sampled delay, referred to as the emission time, which is drawn from a geometric distribution with mean *t*_*d*_. While we assume absorption at the soma and a specific form of signal generation for illustration, the boundary and initial conditions in our framework are flexible and can be adapted to match any relevant experimental setup.

Assuming the above dynamics, we can construct a set of coupled master equations for the time evolution of the probability *P*_*i*_ (*t*) for a particle being at depth level *i* at time *t*, where the hopping and trapping probabilities on the network nodes are taken into account; see [48] for details. The resulting dynamics of individual particles can be described in general by stochastic two-state models [49, 50]. In particular, we previously derived the mean first-passage time (MFPT) of being absorbed in the soma (though for the initial condition of only entering from the dead ends) in terms of the structural parameters {*n, q, p*} by treating the soma as an absorbing boundary [45, 46]. Importantly, we verified that the analytical predictions remain valid for realistic degrees of structural fluctuations (e.g., diversity in the extent of tree along different directions, disorder in the local branching patterns, etc.). Although the high sensitivity of the MFPT to the structural characteristics of dendrites is promising, MFPT is not a directly measurable quantity in dendrites. To realize the practical potential of our approach in technology and medicine, here we extend our proposed formalism and consider the subsequent steps after the particles reach the soma. We assume that each particle emits a temporary pulse after reaching the soma. The accumulation of the pulses generated in many neurons results in an evolving overall signal intensity *I*(*t*) which can be detected externally. *I*(*t*) contains the information of entering, first-passage, and emission times. It indeed reduces to the first-passage time distribution (though shifted by two time steps) in the limit of instantaneous entering and pulse emission, i.e. *t*_*e*_ = *t*_*d*_ = 1. For the general case of *t*_*e*_, *t*_*d*_ *>* 1— where *I*(*t*) deviates from the first-passage time distribution—we demonstrate how the statistical characteristics of *I*(*t*) can be linked to the morphological properties {*n, q, p*} of the dendrite structure. We note that the location of signal generation is in principle arbitrary. Here, we choose the soma as the signal generation point for simplicity—since this choice reduces the problem to an effective 1D lattice—but the approach is extendable to alternative scenarios with different signal generation locations.

The details of calibration of the model parameter *q*, clarification of the required time resolution of measurements, and estimation of the applicability range of our proposed technique are presented in the *Supporting Information* file.

### MAPPING TO REAL DENDRITE STRUCTURES

We first verify the applicability of our mesoscopic approach by mapping the coarse-grained model parameters to the morphological characteristics of real dendrite structures. The depth parameter *n* and node-to-node distance *L* are directly mapped to the extent of the dendritic tree, which primarily depends on the nervous system and neuronal region and type. For instance, cerebellar Purkinje cells of guinea pigs extend up to 200 *μ*m from the soma and have ∼450 dendritic terminals [51]. This corresponds to nearly *n* = 10 generations of junctions which branch out around every *L* = 20 *μ*m.

To map the model parameters *p* and *q* to the structure of real dendrites, we consider the diffusive dynamics of tracer particles along a dendritic tree with protrusions as depicted in Fig. 2(a). The bias *p* in the direction of motion arises from geometrical asymmetries such as tapering of the channel cross-section as well as branching at the junctions into *z*− 1 children (*z* being the number of branches at each junction). The directional bias can be approximated as [48]

**FIG. 2.**
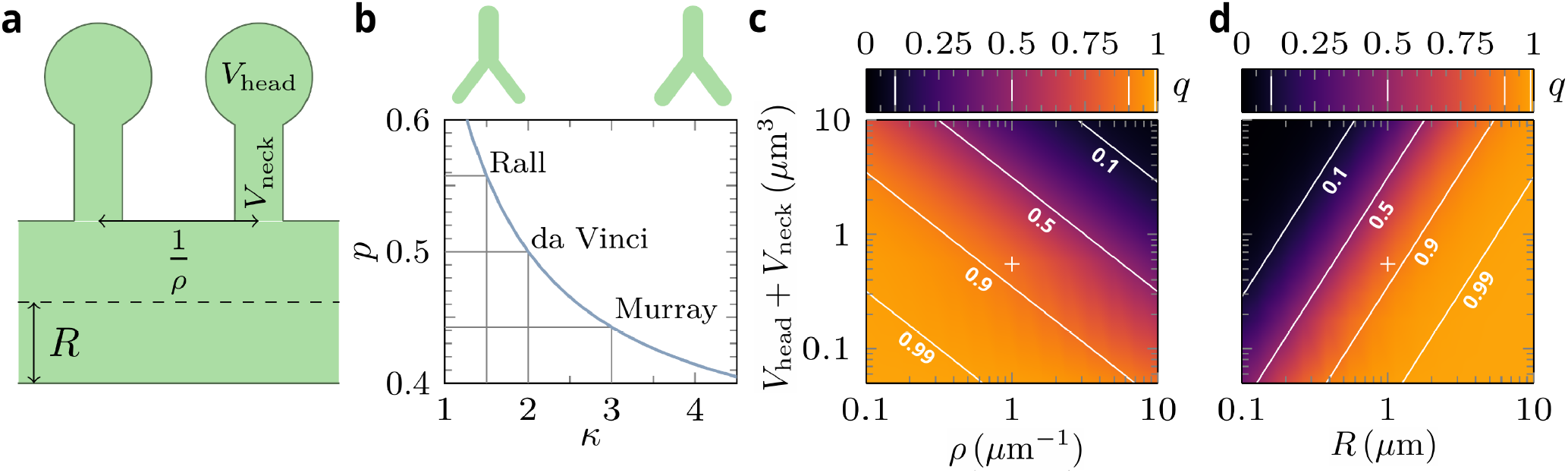
Calibration of the mesoscopic model parameters. (a) Sketch of a section of the dendrite channel. (b) Bias probability *p* in terms of the allometric exponent *κ*. The corresponding points for a couple of known structures are marked. The insets show schematic drawings of channel diameters at different *κ* regimes. (c),(d) Moving probability *q* in the (*V*_head_+*V*_neck_, *ρ*) and (*V*_head_ +*V*_neck_, *R*) planes for the maximum possible time step 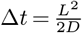. Other parameters: (c) *R* = 1*μ*m, (d) *ρ* = 1*μ*m−1. The solid white lines represent isolines of constant *q* with the indicated values. The crosses mark the set of parameter values (*V*_head_+*V*_neck_ = 0.55 *μ*m3, *ρ* = 1 *μ*m−1, *R* = 1 *μ*m) for a typical healthy dendrite as a reference for comparison.

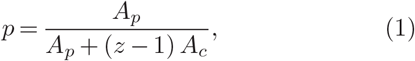

where *A*_*p*_ and *A*_*c*_ denote the cross-sectional areas of the parent and child branches, respectively. The areas can be extracted through the allometric relation 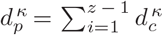 between the diameters *d* and *d* of the parent and child branches at the junction, where *κ* is the allometric exponent [52]. Empirical and theoretical studies suggest representative values of the allometric exponent, with 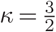 for dendrites of motor neurons [Rall, 1959 [53]], *κ* = 2 for botanical trees [da Vinci’s exponent [54]], and *κ* = 3 for vascular and pulmonary networks [Murray’s exponent[55]]. Other exponents in neuronal context were found to be 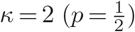 for Purkinje cells, *κ* = 2.5 (*p* = 0.47) for peripheral neuron systems, and *κ* = 3 (*p* = 0.44) for axons [56]. Using the allometric relation, Eq. (1) results in

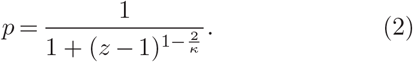

Assuming bifurcations (*z* = 3) and symmetric daughter branches yields, for example, estimated values of *p ≃*0.56, 0.5 and 0.44, for 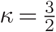, 2, and 3, respectively; see Fig. 2(b). For general channel geometries and driving forces, the bias parameter can be obtained by solving a Fick-Jacobs-like equation [48]. Figure 2(b) illustrates that larger values of *p* correspond to a more pronounced thickening of the channels toward the soma.

To calibrate the probability of motion *q*, we equate the mean time to leave one node in the coarse-grained discrete time model with the mean travel time from the current junction to any of the neighboring ones in the presence of spines. We consider a symmetric branch at which the child channels are connected with equal radii and without leaving a void space. The entrapment of particles inside spines leads to an effective diffusion constant 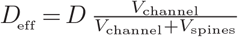 for the diffusive dynamics inside the channel, where *V*_channel_ is the volume of the channel segment, *V*_spines_ is the total volume of spines along it, and *D* is the diffusion constant in the absence of spines [57]. We obtain the following expressions for the moving probability (see *Suppl. Info*. for details):

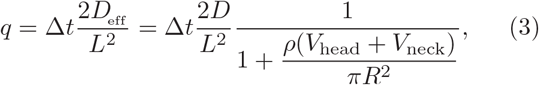

with Δ*t* being the time resolution of measurements, *V*_neck_ and *V*_head_ the spine neck and head volumes, *ρ* the spine density per length unit for regularly spaced spines along the channel, and *R* the radius of the channel segment. The above equation imposes no explicit bound on *q*, however, both *p* and *q* parameters are indeed restricted due to the limited biologically relevant ranges of the structural parameters. For example, *V*_head_+*V*_neck_ ≃0.5 *μ*m3 and *ρ* ≃1 *μ*m−1 represent typical structural parameter values for a healthy dendrite [58]. We also note that the diffusion constant depends on the particle size. Some typical values are: *D* ∼20 (green fluorescent protein (GFP) variants inside spines), ∼100 (large Ca2+ ions inside spines), ∼ 37 (photo activatable GFP, paGFP, inside dendrite channels), and ∼23.5 *μ*m2*/*s (enhanced GFP, eGFP, inside the nucleus) [10, 59, 60]. Using these values, we obtain *t* ≃0.8, 0.5, and 0.2 s for the escape time of eGFP, paGFP, and Ca2+ from spines and *t* ≃8.5, 5.4, and 2.0 s for their travel time from one junction to the next one in a dendritic tree similar to that of cerebellar Purkinje cells but in the absence of spines. The behavior of *q* versus the structural parameters of dendrites is shown in Figs. 2(c),(d). It can be seen that *q* varies monotonically with the structural parameters within their biologically relevant ranges, which allows for mapping of the morphological characteristics of real dendrite structures to the coarse-grained model parameters.

### SIGNAL PROCESSING

To establish a direct link between the dendrite morphology and the statistical characteristics of the detectable signal, our next step is to demonstrate how the coarse-grained model parameters influence the time evolution of the overall signal. We have recently shown that in general the detected signal *I*(*t*) from branched structures develops a peak followed by an exponential decay at long times [48]. The location and height of the peak and the slope of the tail depend on the coarse-grained model parameters and the time scales *t*_*e*_ and *t*_*d*_. For small values of *t*_*e*_ and *t*_*d*_, the signal intensity is nearly equivalent to the first-passage time distribution of passing through the dendritic tree to reach the soma. Note that the signal intensity is invariant under the swapping of *t*_*e*_ and *t*_*d*_, and the asymptotic behavior is governed by the longest time scale. Overall, a faster arrival in the soma and/or a faster emission of the signal is associated with an earlier and higher peak and a steeper tail of *I*(*t*).

To develop a more quantitative understanding of how key model parameters govern the signal intensity evolution, we vary *n, q* and *p* over the biologically relevant ranges and calculate various statistical characteristic of *I*(*t*). Of particular interest is the behavior of the logarithm of the median, 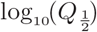. Our previous results revealed that the median of the signal intensity varies monotonically in terms of the structural parameters, even for large values of *t*_*e*_ and *t*_*d*_ [48]. A similar behavior can be observed for the mean or maximum of *I*(*t*). Thus, measuring the median of the signal intensity (or any other quantity in this category) identifies isosurfaces in the (*n, q, p*) space (i.e. a set of admissible {*n, q, p*} values). This is, however, insufficient to uniquely determine these parameters. For a unique determination of the parameters {*n, q, p*}, additional statistical characteristics of *I*(*t*) whose isolines behave differently from the median ones need to be extracted. We tested several quantities, amongst them the variance, skewness, etc. We identified a second category of shape quantities which describe the dispersion of *I*(*t*). The isolines of this category of quantities behave differently from the median’s ones but not significantly from each other. These two independent characteristics of *I*(*t*) allow for identifying at least a one-dimensional manifold in the (*n, q, p*) space. As a representative of the second category, we choose to work with the relative interquartile range Δ*Q*_*r*_ which is a measure of the statistical dispersion of *I*(*t*) and has a smooth behavior upon varying the structural parameters. It is defined as 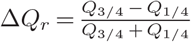, with *Q*_*i/*4_ being the *i*th quartile (for example, the second quartile *Q*_2*/*4_ ≡*Q*_1*/*2_ corresponds to the median). We previously verified that the isolines of Δ*Q*_*r*_ and 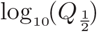 have distinctly different orientations at small *t*_*e*_ and *t*_*d*_ [48]. In this regime, each pair of isolines for a measured set of 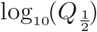 and Δ*Q*_*r*_ intersect at a single point. However, at longer *t*_*e*_ and *t*_*d*_ time scales, the isolines of Δ*Q*_*r*_ may exhibit a nonmonotonic behavior within the relevant range of the structural parameters or become nearly parallel to the isolines of 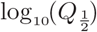 As a result, each pair of isolines for a measured set of 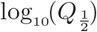and Δ*Q*_*r*_ may have several intersections, meaning that {*n, q, p}* parameters cannot be uniquely determined. We note that identifying the two categories of signal shape parameters may only constrain the coarse-grained model parameters to a 1D manifold in the {*n, q, p}* space. In general, to uniquely determine the structure, either an additional independent quantity can be identified by analyzing other shape parameters of *I*(*t*) or, alternatively, a constitutive relation among the model parameters or the Fourier transform of *I*(*t*)—known as the empirical characteristic function—can be employed. In the following, we assume for simplicity that the two shape parameters, 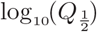 and Δ*Q*_*r*_, suffice to uniquely determine the coarse-grained model parameters.

The sensitivity of the relation between the two sets of structural and signal intensity parameters to the choice of *t*_*e*_ and *t*_*d*_ time scales can be more clearly presented in terms of the degree of information compression when mapping the phase spaces of these two sets to each other. We denote the mapping of structure to signal and vice versa with ψ and ψ−1, respectively. Thus, the connection between the two sets can be represented as 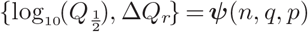 and 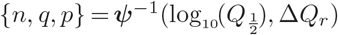 in general. As an example, in (Fig. 3 we show)the mapping of the (*q, n*) plane to the 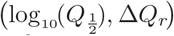 plane for different mean entering *t*_*e*_ and emission *t*_*d*_ times. We regularly sample the phase space of structural parameters (yellow crosses in left panels) and perform extensive simulations to obtain the signal intensity *I*(*t*) and extract its median 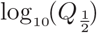 and dispersion Δ*Q*_*r*_ for each sampled set of {*n, q, p*}. For small values of *t*_*e*_ and *t*_*d*_, the structure domain is mapped one-to-one to the signal domain. The mapping consists of a slightly skewed rotation but mapping of two or more distinct points of the {*n, q, p*} parameter space to a same point in the 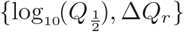space is unlikely, i.e., the map can be inverted. With increasing *t*_*e*_ and *t*_*d*_, the high *q* regions in the structure domain gradually map to highly narrow regions in the signal domain. The compression of information to a space with one less dimension in the limit of large *t*_*e*_ and *t*_*d*_ time scales means that the map cannot be fully inverted anymore. To assess the invertibility limit of *q*, we quantify the compression of the points by the mapping from structure to signal domain; see *Suppl. Info*. for details. By setting a threshold level for the information compression, we can determine the maximum value of *q* (denoted by *q*_max_) up to which the map remains invertible (Suppl. Fig. S1). In Fig. 4(a), *q*_max_ is plotted as a function of *t*_max_ = max(*t*_*e*_, *t*_*d*_), i.e., the longest time scale among the entering and emission times. It reveals that *q*_max_ decays with *t*_max_ roughly as a power-law, which can be presented as 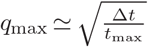 using the fact that *t*_max_ is measured in units of Δ*t*. On the other hand, from Eq. [3] the maximum value of *q* for a given dendritic tree is obtained if spines are absent, leading to 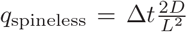 with *D* being the diffusion coefficient in the smooth channel without spines. The full range of *q* is invertible if *q*_spineless_ ≤*q*_max_, which imposes the constraint

**FIG. 3.**
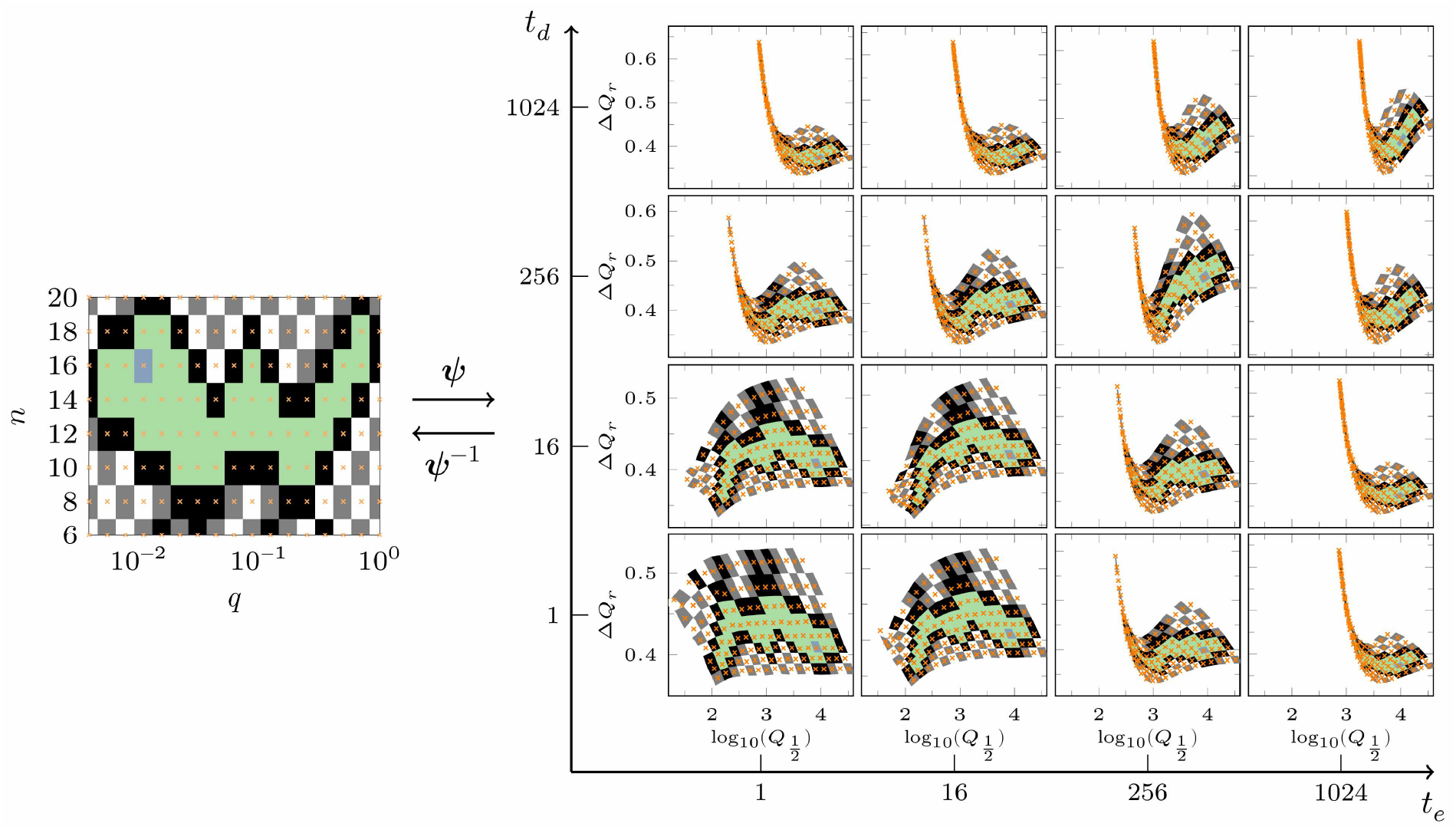
Mapping the structural parameters to the characteristics of the signal intensity (ψ), and vice versa (ψ−1). The mapping of (*q, n*) to 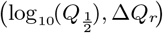 (i.e., the logarithm of the median and the relative interquartile range) is presented for different mean entering *t*_*e*_ and emission *t*_*d*_ times. Other parameter values: *p* = 0.55. For every marked point on the structural parameter domain (yellow crosses), the corresponding location on the signal characteristic domain is extracted numerically. The neighborhood around each pair of connected points are painted with the same color in both domains for clarity.

**FIG. 4.**
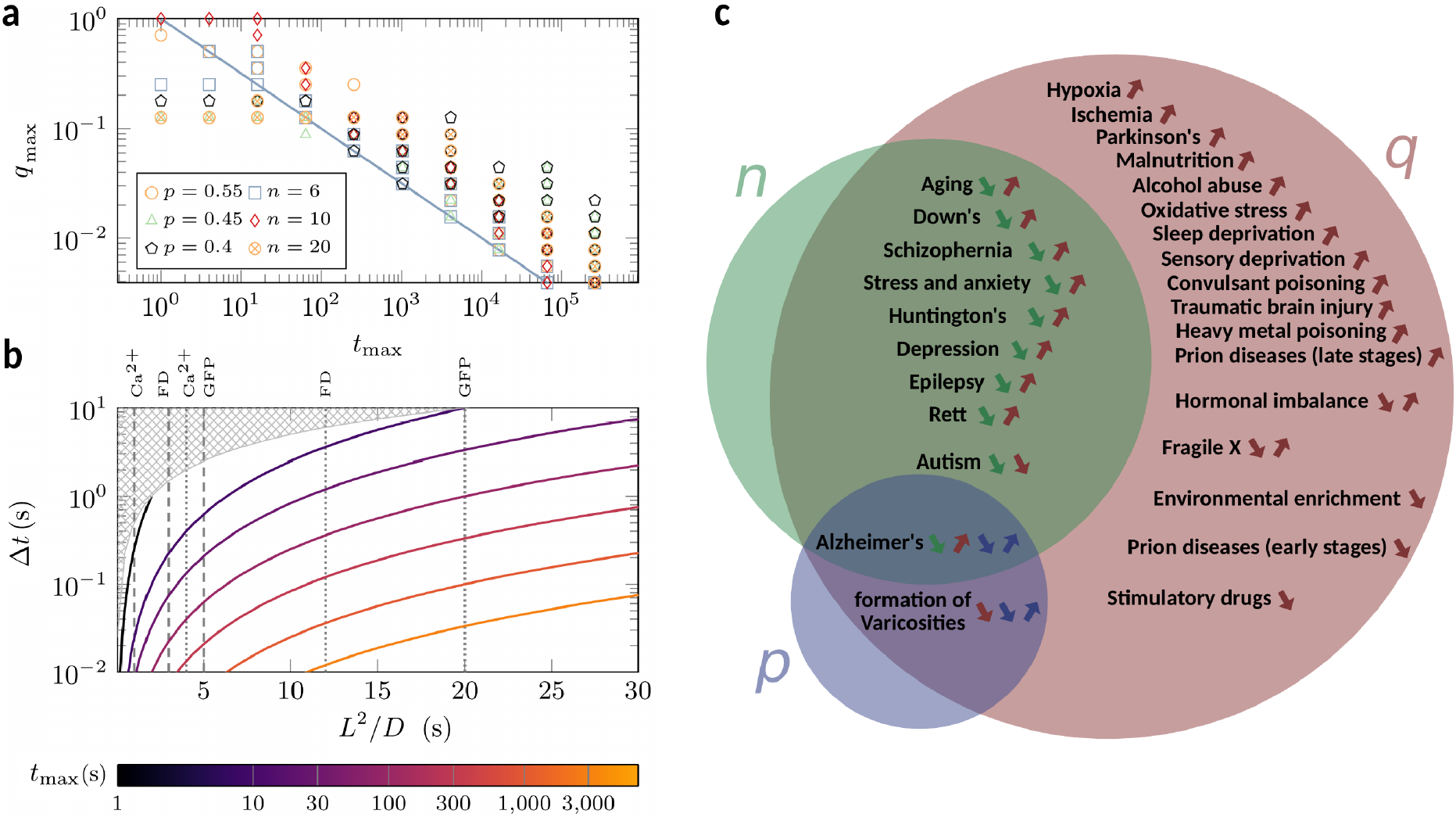
(a) Invertibility threshold *q*_max_ versus the longest time scale *t*_max_= max(*t*_*e*_, *t*_*d*_) for different values of *p* and *n*. The line represents 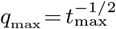 (b) Time resolution of measurements Δ*t* versus diffusive timescale *L*2 */ D* for different values of *t*_max_. The hatched areas indicate the inadmissible region given by Δ*t>L*2*/*2*D*, where the probability *q* would be larger than one. The vertical lines mark the relevant range along the *x*-axis for Ca2+, fluorescein dextran (FD), and green fluorescent protein (GFP) [59, 61]. (c) Pathologies of dendrite morphology, presented in terms of the mesoscopic model parameters *n, p*, and *q* (green, blue, and red colors, respectively). Each arrow indicates the increase or decrease of the corresponding parameter in the course of progression of the given disease. See [17, 62, 63] and references in the main text.

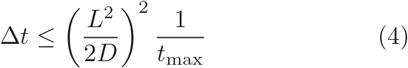

on Δ*t*, as plotted in Fig. 4(b) for different values of *t*_max_. For a given set of dendritic tree structure and tracer particle, the required time resolution of measurements Δ*t* is inversely proportional to *t*_max_. The vertical lines in Fig. 4(b) mark the relevant range of 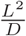 for realistic values of the branching distance *L* and diffusion coefficients *D* for Ca2+, fluorescein dextran (FD), and green fluorescent protein (GFP), as a few examples. It shows that a slower diffusion of tracer particles and shorter entering and emission times lead to a broader possible range for the time resolution of measurements Δ*t*. For time resolutions around Δ*t* ≃0.04 seconds (typical for currently available cameras) and tracer particles with diffusion coefficients similar to GFP, the required entering and emission times can be up to a few minutes. However, in special techniques such as nuclear magnetic resonance spectroscopy, one deals with time resolutions of several seconds, demanding more slowly diffusing particles and shorter entering and emission times on a sub-minute scale.

### MORPHOLOGICAL CHANGES DURING DISEASE PROGRESSION

The morphology of dendrites is broadly affected by aging [19, 21, 32] or neurodegenerative disorders such as Alzheimer’s disease [19, 20, 22–24], autism spectrum disorders [19, 27–30], epilepsy disorder [64], schizophrenia [19, 25, 26], Down’s syndrome [65], fragile X syndrome [30, 31], prion diseases [66], and stress-related disorders [67]. The affected morphological properties include the overall extent of dendritic tree, the segmental increase of dendrite diameter towards the soma, the population of branches, the thickness and curvature of dendrite shafts, and the morphology, density, and spatial distribution of spines. Here we clarify how these morphological changes influence our mesoscopic model parameters {*n, q, p }*. This information is then used in Fig. 4(c) to categorize the neurodegenerative disorders—those for which clear trends for the pathological changes of dendrite structure have been reported in the literature—based on the expected trends of the model parameters in the course of disease progression.

Variation of the extent of dendritic tree trivially influences the depth parameter *n*. Reduction of the tree extent is the observed trend during aging and several disorders such as Down’s syndrome, schizophrenia, Alzheimer’s disease, autism spectrum disorders, epilepsy disorder, stress-related disorders, Huntington’s disease, etc. Increasing of the tree extent due to neurodegenerative disorders has not been reported to our knowledge.

The moving probability *q*, given by Eq. (3), is the only parameter affected by the presence of spines. *q* increases with decreasing spine size or density as observed, e.g., in aging, Down’s syndrome, Alzheimer’s disease, and schizophrenia; see Fig. 4(c) for an extended list of relevant disorders. Conversely, the spine density increases in a few cases such as autism spectrum disorders, fragile X syndrome, and hormonal imbalance, leading to the decrease of *q*. Nevertheless, the pathology of fragile X makes the prediction of *q* variations complicated: The increase of spine density (decrease of *q*) is accompanied by the shrinkage of spine size (increase of *q*); thus, the two effects compete and may even compensate each other such that *q* remains unchanged. The influence of hormonal imbalance on *q* depends on the hormone type and whether there is a deficiency or surplus.

The disorders that change the width of dendrite shafts influence the bias parameter *p* in general. Particularly, the decrease in dendritic arborization is often correlated with the overall reduction of the channel width. The details of width reduction determine the direction of changes of *p*: While a uniform reduction of channel radii effectively increases *p* (and decreases *q*), a radius-dependent reduction may change *p* in both directions. Moreover, an inhomogeneous spatial pattern of *p* can be caused by local changes of the channel width. For instance, local thinning of channel occurs in the vicinity of amyloid plaque deposition in Alzheimer’s disease. Finally, the disorders that reduce the population of branches can increase the average node-to-node distance *L*, resulting in smaller *q* and *n*.

We note that the pathology of spine and dendrite structure is more complicated in other diseases. For instance, distortion of spine shape observed in most mental retardations makes the prediction of the trend of *q* difficult. Despite the currently available information discussed above, there is a lack of quantitative studies to clarify the impact of various neurodegenerative disorders on dendritic spine, tree metric, and topological morphology.

## DISCUSSION

We have proven that the parameters {*n, q, p}* of our mesoscopic model can be extracted by analyzing the detectable temporarily signal generated by a large population of neurons, provided that the time scales of entering the dendrites and emission of signal after reaching the soma are sufficiently small compared to the mean first-passage time of passing the dendritic tree. Although we constrained our analysis to signals formed by spontaneous pulses emitted by the particles in the soma with activation probability 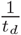, signals of other forms can be easily obtained from the signal studied in the present work. For example, if one seeks insight into the ability of neurons to integrate spine-derived (concentration) signals, the number of particles in the soma that have not yet emitted their pulse (i.e. are still active in this case) would be of interest. This quantity at time *t* is given by *t*_*d*_ *I*(*t*+1), i.e. our measured signal shifted by one time step to the left and scaled by the mean activation time. In Suppl. Fig. S3, we present this signal alongside the time evolution of the fraction of particles in the soma that have not yet emitted their pulse for different values of *t*_*e*_ and *t*_*d*_ and for healthy versus differently degenerated dendritic trees.

On the other hand, the model parameters {*n, q, p}* can be directly linked to the morphology of real dendrites via Eqs. (2) and (3). Since there are several morphological characteristics on the right hand sides of these equations, they cannot be uniquely determined by a given set of {*n, q, p}*. Nevertheless, most neurodegenerative disorders affect only a few of the morphological properties of dendrites. Therefore, by conducting regular patient monitoring for a given disease, the observed changes in the parameters {*n, q, p}* can be attributed to the changes of the morphological properties relevant to that specific disease. For instance, the growth rate of *q* and reduction rate of *n* for a patient with schizophrenia reflect, respectively, how fast the mean spine volume and the extent of dendritic tree are shrinking over time.

To link the detected signal intensity to the mesoscopic model parameters we have considered an ideal regular tree structure, while real dendritic trees are irregularly branched, spines have diverse sizes, and their spatial distribution is inhomogeneous. These fluctuations naturally cause variations in the corresponding model parameters {*n, q, p}*. However, we verified in our previous study [45] that the analytical results for the first-passage times of passing a regular tree structure remain valid when realistic degrees of global fluctuations of the structural parameters across the tree or local structural irregularities in the branching patterns are considered. Since the dependence of the signal intensity on the dendrite morphology is due to the contribution of the fist-passage times (and not the entering *t*_*e*_ and emission *t*_*d*_ times), we conclude that the presented results in the current study remain valid under typical structural irregularities and fluctuations observed in real neuronal dendrites.

We have characterized the behavior of the signal intensity *I*(*t*) by two quantities, the logarithm of the median 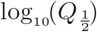 and the relative interquartile range Δ*Q*_*r*_. The former is a representative of a category of the statistical measures including the mean, median, and maximum of *I*(*t*). The latter quantifies the statistical dispersion of *I*(*t*) and behaves similar to quantities such as the normalized variance and skewness. One may still identify further independent quantities by analyzing other moments of *I*(*t*). Additional statistical measures of *I*(*t*)—which vary smoothly with the parameters {*n, q, p*} and exhibit isolines that differ from those of 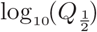and Δ*Q*_*r*_—can in principle improve the accuracy of the extracted values of the model parameters {*n, q, p*}.

Our study focuses on solving the technical problem of processing the generated signal and linking it to the dendrite morphology. Nevertheless, preliminary steps should first be taken to generate the signal. This includes the transport of cargos to desired regions of brain tissue, entering the dendrites, travelling to the soma, and generating a temporary signal. Our proposed measurement procedure is summarized in Fig. 5(a). While addressing the technical issues of each preliminary step is beyond the scope of this study, obtaining a detectable signal is indeed feasible using the currently available technologies. In the following, we discuss possible methods for each of the measurement procedure steps:

**FIG. 5.**
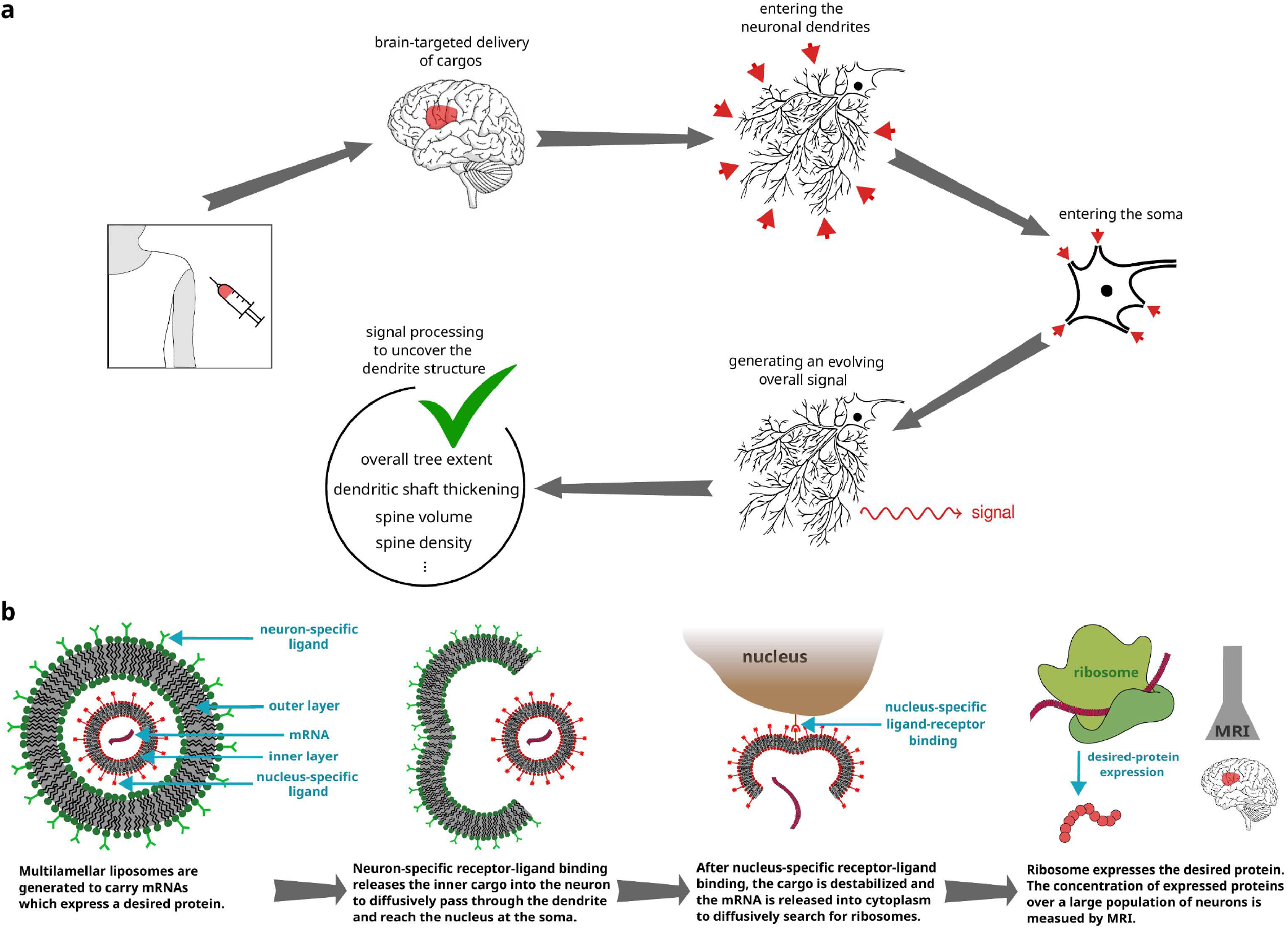
(a) Schematic of our proposed noninvasive technique for a fast indirect measurement of the structural properties of dendrites based on processing a signal generated by a large population of neurons. By conducting regular measurements for a given patient, essential information about the morphological evolution of dendrites in the course of neurodegenerative disease or treatment progression can be extracted. (b) Schematic illustration of multilamellar liposomes designed to generate a specific protein. As a detectable temporary signal, the concentration of the expressed protein across the brain region of interest is externally monitored by the MRI technique.

1. **Transport of cargos to desired regions of brain tissue by means of brain-targeted drug delivery techniques**. Promising strategies have been developed so far to deliver drugs specifically to the brain to treat neurological disorders while minimizing systemic side effects [68–70]. Some of the currently feasible techniques include nanoparticle-based delivery [68, 71–78], focused ultrasound (non-invasive technique which offers spatially targeted drug delivery by transiently disrupting the blood-brain barrier (BBB) to pass through) [79], and carrier-mediated transport (utilizing endogenous transport systems like glucose or amino acid transporters facilitates drug transport across the BBB. Drug molecules are conjugated with ligands that target these transporters to enhance brain uptake) [80]. Moreover, injection into the spinal chord fluid or into ventricles could be an option to pass the BBB as well.
2. **Entering the neuronal dendrites and passing through their complex structure to reach the soma**. After reaching the area around the neurons, the contents of the cargos can enter the neuron by means of neuron-specific receptor-ligand binding [73–78]. This would mainly occur through the dendritic tree rather than axon or soma since the outer area of the neuron is mainly formed by the dendritic tree. Nevertheless, the contribution of entering from soma or axon to the generated signal can be evaluated and subtracted, as long as the axons and somata do not undergo morphological changes in the course of disease progression or treatment.
3. **Generating a temporary signal and detecting it**. Here, we mean any kind of detectable signal such as, but not limited to, electric or magnetic fields generated by many neurons. There are powerful noninvasive techniques for real-time tracking of brain activities. Electro- and magneto-encephalography for electric and magnetic field detection are established neurotherapeutic tools [39, 40]. Another possibility is to employ nuclear magnetic resonance spectroscopy (NMRS), which allows for noninvasive measurements of the concentration of different neuro-chemicals within a volume of brain down to a few cubic centimeters [41, 42]. The concentrations of substances generated in the somata of neurons can be obtained via NMRS with a time resolution of a few seconds which, depending on the diffusion constant of the particles, can remain within the feasibility range of our proposed method [81]. Positron emission tomography and magnetic resonance imaging (MRI) can be also employed to measure the concentration of neurochemicals [43, 44].
4. **Processing the evolving overall signal**. After detecting the signal, the approach developed in this paper enables one to process the signal intensity to uncover the morphology of dendrites.

As a more detailed plan for generating a detectable signal, we propose a protein expression scenario by injecting specific mRNAs carried by multilamellar liposomes; see Fig. 5(b). The concept of producing multilamellar liposomes is currently feasible and has been realized in the context of cell activity regulation, immunotherapy, and vaccination [82–84]. Transport of liposomes to desired regions of brain and uptake of them by neurons have been feasible by modifying their surface with ligands targeting specific receptors on brain endothelial cells or neuronal dendrites [73–78]. Upon neuron-specific receptor-ligand binding, the multilamellar liposome enters the dendrite and loses its outer layer, leading to the release of the inner cargo into the cytoplasm. To enhance the dendrite specific entering, the endocytosis events around the synapses can be harnessed. For example, there is evidence that AMPA receptors are preferentially endocytosed around synapses [85]. The inner cargo is conjugated with nucleus-specific ligands [86–88] and diffuses inside the dendritic tree until it enters the soma and reaches the nucleus. The cargo can be designed to be destabilized or dissolved after the nucleus-specific receptor-ligand binding pins it to the exterior of the nucleus. This can be achieved, e.g., through specific proteolytic enzymes or pH-sensitive components in the cargo structure or adjusting the concentration of aqueous ionic solutions inside the cargo [89] to respond to the environmental differences between the region around the nucleus and the rest of the cytoplasm. The destabilized cargo releases mRNAs into the cytoplasm, which will diffusively search for ribosomes to produce a desired protein, such as ferritin. A typical neuron contains millions of ribosomes but their homeostatic distribution is still unknown [90, 91] (though recent studies revealed spacial inhomogeneities in the protein translation across the neuron, attributed to the spacial distribution of mRNAs and potential local specialization of ribosomes [91, 92]). The expressed protein should be harmless and degrade over a reasonable time. Variations of the concentration of this protein over a large population of cells can be externally detected. For example, expression of ferritin can be monitored by the MRI technique [44]. We note that the presented formalism in the previous sections to obtain the first-passage time distribution of reaching the soma can be straightforwardly extended to calculate the additional first-passage time distribution of reaching from the soma to the distributed ribosomes. Moreover, here we considered a spontaneous signal emission but the formalism can be adapted to other scenarios such as a gradually degrading signal.

The clinical translation of these proposed techniques certainly requires rigorous testing and validation to ensure the safety and specificity of the novel approaches. Uptake and transport of cargos can be initially tested, e.g., in cultured murine neurons. For the plan proposed in Fig. 5(b) based on the expression of proteins by ribosomes, spatial distribution of ribosomes in different types of healthy neurons needs to be determined. We expect that our proposals can potentially trigger active research and development in the fields of neuroscience, molecular engineering, and pharmacology.

To conclude, a framework has been developed to link the statistical characteristics of a detectable signal generated after reaching the somata of neurons to the morphological properties of neuronal dendrite structures. Our results open the possibility of indirectly monitoring the morphological evolution of dendrites in the course of neurodegenerative disorder progression. The mesoscopic approach presented in this study can be generalized to cope with further details of transport in real dendrite structures, such as handling the memory effects and aging inside spines [93] or to include active transport of cargos along microtubules [94]. Besides drawing conclusions regarding the morphological changes of dendrite structures, investigation of the first-passage properties of stochastic motion inside dendrites can deliver vital information about the ability to preserve local concentrations or induce concentration gradients of ions and molecules. These are tightly connected to neural functions and allow for drawing important physiological conclusions. The proposed approach also provides a route into a variety of other stochastic transport phenomena e.g. in varying energy landscapes, branched macromolecules and polymers, and labyrinthine environments with absorbing boundaries.

## Supplementary Information to Tracking the Morphological Evolution of Neuronal Dendrites by First-Passage Analysis

### Mapping dendritic structure to the model parameter *q*

The mean escape time ⟨*t⟩* from a junction to any neighboring furcation can be expressed, on the one hand, in terms of channel geometry as 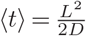, assuming diffusive dynamics. On the other hand, using the discrete-time framework of the model with observation time resolution Δ*t*, it is given by 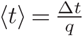. Equating these two expressions yields a relation between the moving probability *q* and the geometric parameters of the dendritic structure, 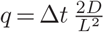. However, we have assumed a smooth channel so far, thus, this relation does not yet account for the effects of trapping in dendritic spines. In the following, we show how these effects can be incorporated into the framework.

We note that transient trapping events along the channel do not induce any bias in the motion towards one end of the channel segment, thus, no modification is required in the calibration relation for the parameter *p*. However, for the escape time (*t*), frequent interruption of motion by entrapment events in spines has a considerable impact. To keep the model traceable, this impact is taken into account by an effective asymptotic diffusion constant *D*_eff_. Previous studies have already calculated such an effective diffusion constant in a geometry almost tailored to the diffusive transport in spiny dendrites [57]. The geometry considered there consists of a cylindrical tube from which identical spines protrude periodically. The spines were modeled as spherical cavities connected to the main shaft by narrow cylindrical necks. The effective diffusion constant derived in [57], adapted to our application, is given by

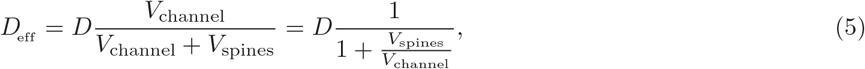

where *D* is the diffusion constant without protrusions, *V*_channel_ the channel volume, and *V*_spines_ the total volume of spines. Let us assume a uniform distribution of spines along the dendritic channel with the density *ρ* per length unit. Denoting the spine head volume with *V*_head_ and neck volume with *V*_neck_, the ratio 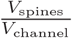can be written as

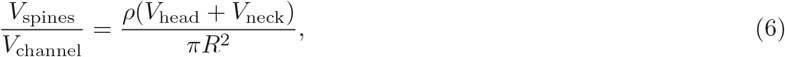

where *R* is the channel radius. Substituting *D*_eff_ into the calibration relation for *q* yields

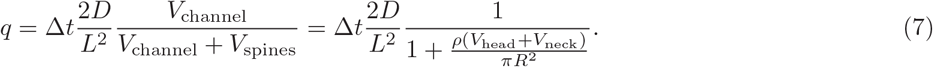

As *q* is a probability, it cannot be larger than one, imposing a constraint on the time resolution of observation Δ*t*. Since the relation 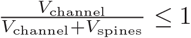 always holds, the condition

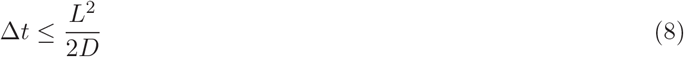

ensures that *q* always remains as a valid probability, i.e., *q* ≤ 1.

To derive the above calibration relation, we have made a few simplifying assumptions for the diffusive dynamics of tracer particles inside dendritic channels. For example, a constant distance *L* between successive junctions is assumed. However, *L* may vary in real dendrite structures, not only between the segments within one generation but also between different generations. The primary segments of apical dendrites of pyramidal neurons and terminal segments of all dendrites are reported to be longer than intermediate segments. As a result, *q* should slightly vary with the depth of dendritic tree due to its *L*-dependence.

Additionally, spines are inhomogeneously distributed over the dendritic tree. There are almost no spines very close to the soma but the spine number density rapidly grows and saturates after a short distance from the soma. However, the gradual thinning of the channel towards dendritic terminals practically increases the trapping probability inside spines and, hence, decreases *q*.

Nevertheless, we verified in our previous study [45] that the analytical results for the first-passage times of passing a regular tree structure remain valid when realistic degrees of global fluctuations of the structural parameters across the tree or local structural irregularities in the local branching patterns are considered. To conclude, the structural parameters which enter into the calibration relations for *q* and *p* parameters should be considered as average values over the entire dendritic tree.

### CONSIDERATIONS FOR CHOOSING THE TIME RESOLUTION OF MEASUREMENTS

In this section we discuss the choice of the time resolution of measurements Δ*t* and provide a rough estimate of the applicability range of our proposed method. Figure 3 of the manuscript revealed that the invertibility of mapping the structural parameters to the signal characteristics breaks down for large entering *t*_*e*_ and/or emission *t*_*d*_ times. We further clarified that only the mapping of high *q* regions is problematic, while low *q* regions can be resolved even for very large values of *t*_*e*_ and *t*_*d*_. To assess the invertibility limit of *q*, in the following we quantify the compression of the points by the mapping. For any given point in the structural parameter space and the corresponding point in the signal domain, the degree of compression is determined by calculating two distances: the minimum distance *ℓ*_struct_ between the selected point and all other points in the struct(ural parameter)space and the minimum distance *ℓ*_signal_ between the corresponding point and all other points in the 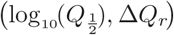plane of signal characteristics. In each of the two domains, the distance between a pair of points (*x*_1_, *y*_1_) and (*x*_2_, *y*_2_) is calculated using the metric

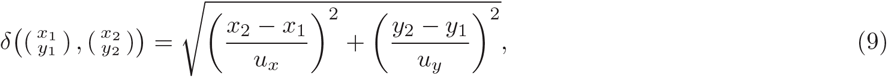

where (*x*_*i*_, *y*_*i*_) can be any pair of the structural parameters {*p, q, n*} or a point in the 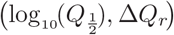 plane of signal characteristics. *u*_*x*_ and *u*_*y*_ denote the total variation range along *x* and *y* axes, respectively. Note that *u*_*x*_ and *u*_*y*_ in the signal domain are determined by combining the variation range of 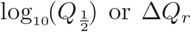 over all choices of the entering and emission times.

From the minimum distances *ℓ*_struct_ and *ℓ*_signal_, the volumes of the neighbourhoods in the two domains can be estimated as 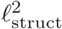 and 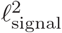, respectively. We introduce the ratio of the volumes 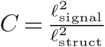 (hereafter refereed to as compression ratio) as a measure of the degree of mapping compression. An invertible mapping requires *C >* 0. In this regard, the compression ratio has similarities with the Jacobian determinant of the mapping. The difference between them is that the distances to all other points in each domain are taken into account in *C* whereas for the Jacobian only the neighborhood in each domain enters. Therefore, *C* is a stronger measure for invertibility because it vanishes even when a point far away from the one where *C* is calculated is mapped to the same point in the signal domain, but the Jacobian determinant cannot capture it due to its local nature. In the case of a globally invertible map, *C* is an estimate for the absolute value of the Jacobian determinant.

Setting a lower compression threshold *C*_min_ allows us to identify the points for which the invertibility of the map between structural and signal domains practically breaks down. By choosing a threshold value *C*_min_ = 0.05, we identify the points where mapping the structural parameters to the signal characteristics plane is not invertible. This procedure is visualized in Fig. S1 for a constant *p* and various entering and emission times. Next, we determine *q*_max_ as the maximum value of *q* up to which the map is invertible for all values of the other dimension of structural parameters (*n* in the cases presented in Fig. S1). We checked that the choice of the threshold level *C*_min_ has no qualitative impact on the behavior of *q*_max_ and only induces minor quantitative changes.

In Fig. S2, *q*_max_ is plotted as a function of the entering time *t*_*e*_ and the emission time *t*_*d*_ for a given value of *p*. It can be seen that the isolines of constant *q*_max_ are roughly square-shaped which evidences that *q*_max_ is a function of the largest time scale, i.e. *t*_max_ = max(*t*_*e*_, *t*_*d*_). This is confirmed in Fig. 4(a) of the manuscript, where *q*_max_ is plotted as a function of *t*_max_ for different values of the structural parameters. The overall trend of *q*_max_ can be roughly captured by a power-law scaling 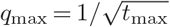. The observed deviations from the power-law scaling originate from the intrinsic stochasticities as well as our method of compression ratio calculation. For small values of *t*_*e*_, *t*_*d*_ and *p* and large values of *n*, the points of the structural parameters domain are mapped onto a patch with low but nonzero extension along the Δ*Q*_*r*_ direction in the signal domain. By increasing the entering or emission time, the points collapse on a nonmonotonic curve which covers a much larger range along Δ*Q*_*r*_ direction; see Fig. S1(b). This results in larger *u*_*x*_ or *u*_*y*_ in the metric Eq. [9], thus, smaller *C* values. Hence, those points may be considered as non-invertible despite that their mapping to the signal domain is properly resolved.

From the power-law relation between *q*_max_ and *t*_max_ and the fact that the time scales *t*_*e*_ and *t*_*d*_ are measured in units of the time resolution of observation Δ*t*, it reads

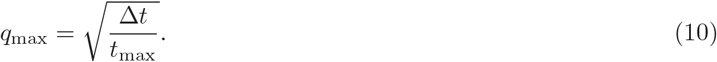

According to Eq. [7], the value of *q* for a given dendritic tree can be almost arbitrarily tuned through Δ*t*. If the upper estimate of *q* from Eq. [7] (obtained for a smooth channel, i.e. *V*_spines_=0) is less than *q*_max_ given by Eq. [10], variations of *q* due to morphological changes of spines can be fully resolved with our proposed approach. By equating Eq. [10] with Eq. [7] at *V*_spines_=0 we obtain the relation

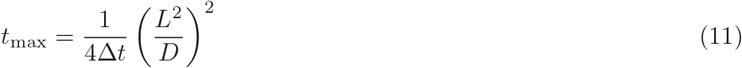

between *t*_max_= max(*t*, *t*), the time resolution Δ*t*, and the diffusive timescale 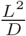 (*D* is the diffusion coefficient of the tracer particles in the spineless dendritic channel). Therefore, for a given set of dendritic tree structure and tracer particle, the required time resolution of measurements Δ*t* is inversely proportional to *t*_max_, i.e. the maximum time scale among the entering and emission times *t*_*e*_ and *t*_*d*_. In Fig. 4(b) of the manuscript, Δ*t* is plotted versus the diffusive timescale 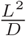 for different values of *t*_max._ The vertical lines mark the relevant range of 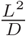 for realistic values of the branching distance *L* and diffusion coefficients *D* for Ca2+, fluorescein dextran (FD), and green fluorescent protein (GFP), as a few examples. It shows that a slower diffusion of tracer particles and shorter entering and emission times lead to a broader possible range for the time resolution of measurements Δ*t*. For time resolutions around Δ*t* ≃ 0.04 s (typical for currently available cameras) and tracer particles with diffusion coefficients similar to GFP, the required entering and emission times can be up to a few minutes. However, in special techniques such as nuclear magnetic resonance spectroscopy, one deals with time resolutions of several seconds, demanding more slowly diffusing particles and shorter entering and emission times on a sub-minute scale.

**Suppl. Fig. S1:**
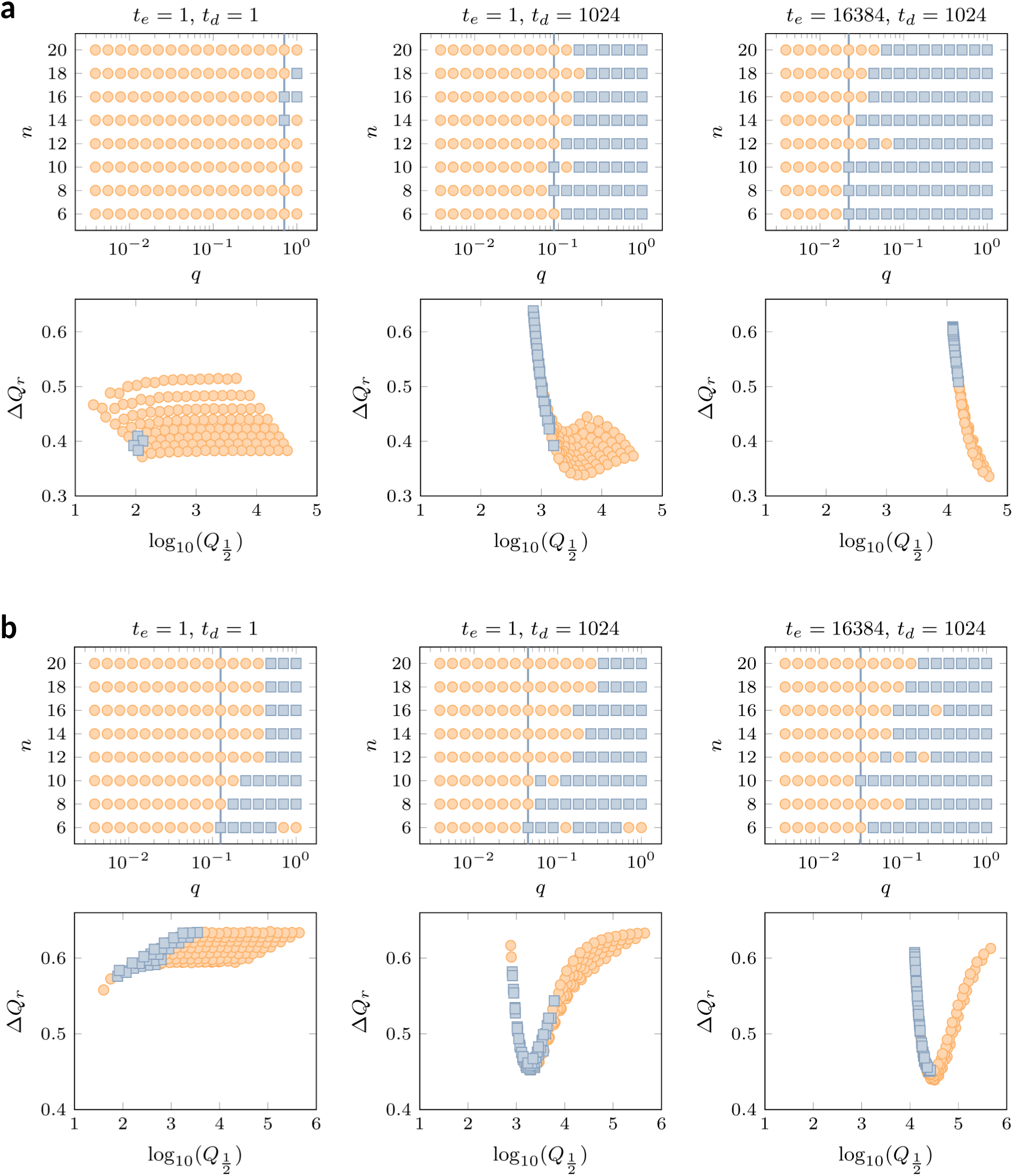
Visualisation of mapping the structural parameters (*q, n*) to the signal characteristics 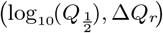 for different entering times *t*_*e*_ and emission times *t*_*d*_. Other parameter values: (a) *p* = 0.55, (b) *p* = 0.45. The points where the map is invertible (not invertible) for the compression threshold *C* = 0.05 are shown with orange circles (blue squares). The blue vertical line marks *q*_max_, up to which the map is invertible for all values of *n*.

**Suppl. Fig. S2:**
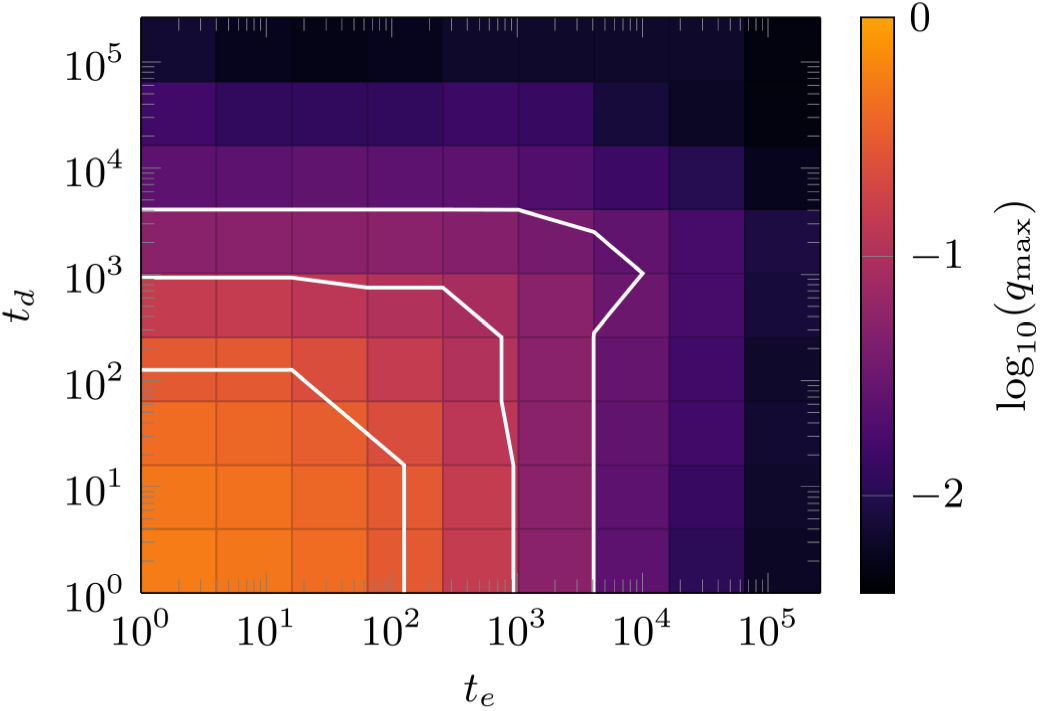
Logarithm of *q*_max_ (i.e. the maximum value of *q* up to which the map from (*q, n*) to signal domain is invertible) as a function of entering and emission times *t*_*e*_ and *t*_*d*_. *q*_max_ is extracted for *p* = 0.55 and *C*_min_ = 0.05. The solid white lines represent isolines of constant *q*_max_.

**Suppl. Fig. S3:**
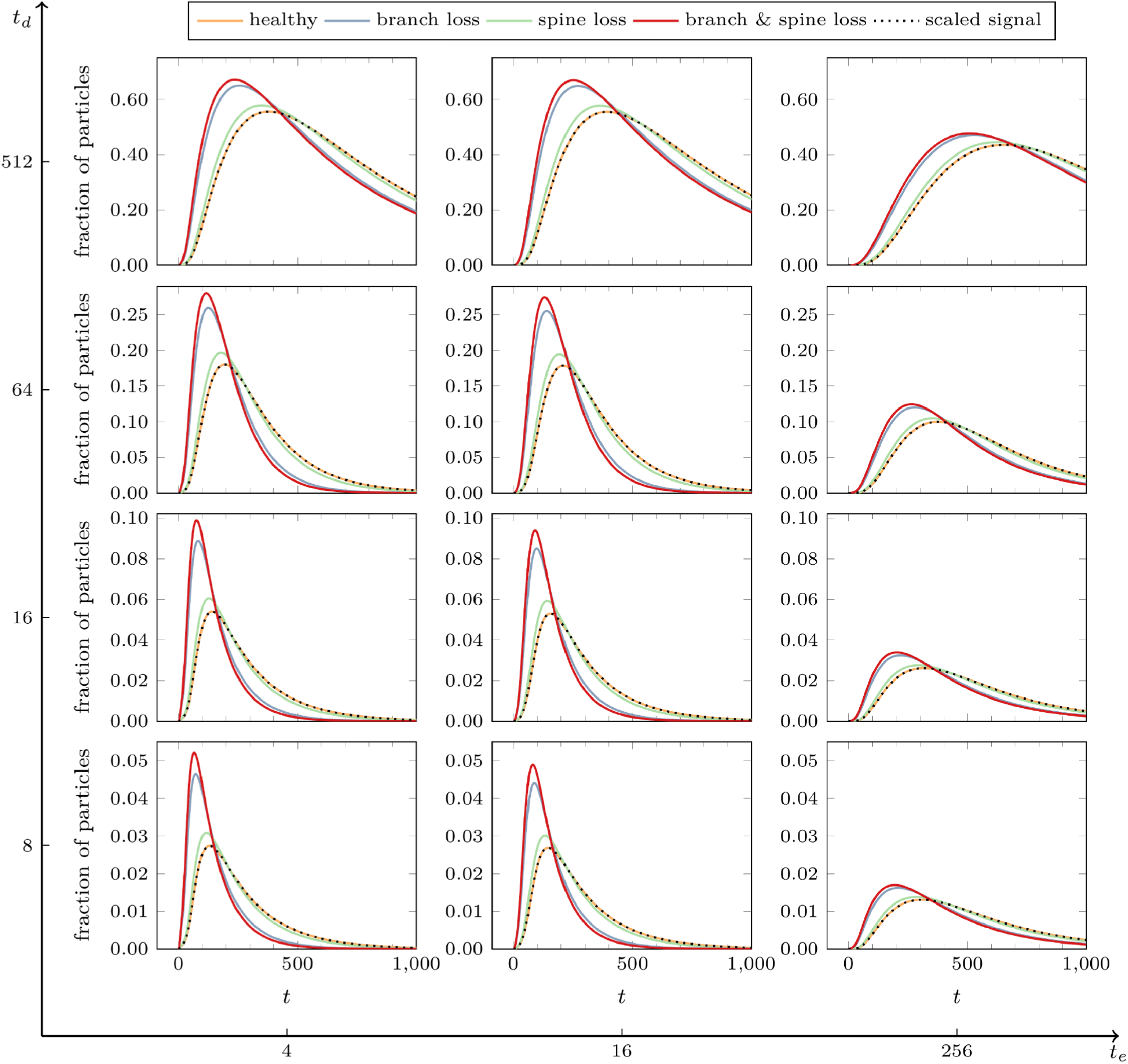
Fraction of particles in the soma as a function of time for healthy and differently degenerated dendritic trees for different entering and emission times *t*_*e*_ and *t*_*d*_. For the healthy dendrite (orange line) the following parameters were assumed: *n* = 10, *ρ* = 1 *μ*m^−1^, *V*_head_ + *V*_neck_ = 0.55 *μ*m3 and *R* = 1 *μ*m. The time resolution was chosen to be 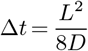 corresponding to Δ*t* = 2.5 s for a dendrite with mean branch length *L* = 20 *μ*m and particles with a diffusion constant *D* similar to GFP. The degeneracies were branch loss (blue line) where the tree has lost three generations of branches, spine loss (green line) where the dendrites have lost three quarters of their spine volume as well as the combination of both (red line). Increasing *t*_*d*_ increases the fraction of particles in the soma leading to a broader and higher curve. Increasing *t*_*e*_, on the other hand, leads to broader and flatter curves because of the restricted influx of particles. For the healthy dendrite, the signal *I/*N (generated by the accumulated pulses of the particles in the soma scaled by *t*_*d*_ and shifted by one step to the left) is shown with the dotted line. This line coincides with the one for particle fraction, exemplifying that the particle fraction in the soma can be obtained by the signal and vice versa.

